# Reinforcement meta-learning optimizes visuomotor learning

**DOI:** 10.1101/2020.01.19.912048

**Authors:** Taisei Sugiyama, Nicolas Schweighofer, Jun Izawa

**Affiliations:** Empowerment Informatics, University of Tsukuba, Tsukuba, Ibaraki, 305-8573, Japan; Biokinesiology and Physical Therapy, University of Southern California, Los Angeles, CA, 90089-9006 USA; Engineering, Information, and Systems, University of Tsukuba, Ibaraki 305-8573, Japan

**Keywords:** meta-learning, reinforcement learning, motor learning, cerebellum, basal ganglia, reward, punishment, saving

## Abstract

Reinforcement learning enables the brain to learn optimal action selection, such as go or not go, by forming state-action and action-outcome associations. Does this mechanism also optimize the brain’s willingness to learn, such as learn or not learn? Learning to learn by rewards, i.e., reinforcement meta-learning, is a crucial mechanism for machines to develop flexibility in learning, which is also considered in the brain without empirical examinations. Here, we show that humans learn to learn or not learn to maximize rewards in visuomotor learning tasks. We also show that this regulation of learning is not a motivational bias but is a result of an instrumental, active process, which takes into account the learning-outcome structure. Our results thus demonstrate the existence of reinforcement meta-learning in the human brain. Because motor learning is a process of minimizing sensory errors, our findings uncover an essential mechanism of interaction between reward and error.

---

Learning is considered a skill that can be improved by experience and motivation^1,2^. In fact, animals and humans often exhibit accelerated learning over training sessions^3^. How does the brain learn to learn quickly in variable tasks and environments?

Accelerated learning has often been reported in motor learning tasks where human participants learn to compensate for force or visual perturbations to generate a planned movement trajectory^4–8^. After motor memories formed in the initial learning session have been washed out, learning in the second learning session becomes faster than learning in the first session, a phenomenon that is known as the ‘saving effect’^4,9^. In this task, motor learning is considered a process of minimizing sensory prediction errors, i.e., the discrepancy between the generated and predicted movement trajectories is independent of the motivational signals^10–12^. Research has shown that this saving effect is achieved by updating the learning rate (i.e., the policy of how much the motor memory is updated in response to the perceived sensory errors), which is driven by prior experience of the errors^7^. Additionally, recent studies suggest that there is a significant effect of motivational signals on the learning speed^13–17^. How does the brain incorporate motivational signals into the history of errors to regulate learning rates?

One suggested mechanism is a passive, Pavlovian (cue-outcome-based) process in which the valence (reward or punishment) biases the learning rates. For example, in the go and no-go learning task, valence biases the speeds as well as the asymptotes of learning curves^18^. In saccadic adaptation, rewards increase the speeds of adaptation, which is induced by dopaminergic modulation of error signals^19^. While this motivational, hard-wired process might induce biases in error sensitivity of motor learning, it might not explain variations in reward influence of motor learning: some studies show that learning is facilitated by motivational signals^14^, while others report no effect^16,17^, or even decelerated learning^15^. Thus, these variations imply that another mechanism other than the hard-wired mechanism regulates the speed of learning.

In theory, the ultimate goal of motor learning is to maximize future rewards^20^. Thus, reinforcement learning, which is driven only by reward feedback without any sensory prediction error, also forms motor memory^21–23^. Because the spatial generalization function of learned memory, which reflects properties of neural basis of adaptation, is significantly different for error-based and reward-based learning, two dissociable neural mechanisms are likely involved in error- and reward-based motor learning^24^.

However, how these two learning systems interact with each other is still not known. Here, we hypothesized that the integration of error-based and reward-based learning systems reinforces motor learning, which we call ‘reinforcement meta-learning.’ In contrast to a passive process, this learning process is active and instrumental (action-outcome-based), where a higher-level reinforcement learning mechanism trains a lower-level motor learning mechanism.

This idea of reinforcement meta-learning is based on a theory developed in machine learning where the parameters characterizing learning behaviors such as learning rates are modulated by high-order reinforcement learning^25,26^. This idea can account for changes in learning rates in decision-making tasks in animals and humans^27^. However, the existence of reinforcement meta-learning in the brain has been difficult to verify experimentally because in decision-making tasks, both learning and meta-learning are updated by a reward feedback, obscuring the contribution of each learning layer.

Here, we devise a reinforcement meta-learning task in which the feedback for learning and that for meta-learning are dissociated: sensory error feedback is provided in a motor learning trial, and reward feedback that depends on the rate of motor learning is provided in a subsequent meta-learning trial. If the brain employs reinforcement meta-learning, it integrates sensory error feedback and reward feedback to form associations between these two to regulate how large motor commands should be updated in response to the observed sensory error. Thus, manipulating the relationship between the extent of learning and the reward feedback should influence how motor learning rates are modulated as a result of meta-learning regardless of which reward and punishment are presented. Alternatively, if the modulation of learning rate is hard-wired to valence, such manipulation might not influence learning rates.

## Results

Forty one healthy participants gave informed consent before participating in the experiment, which was approved by the Institutional Review Board at the University of Tsukuba. One subject was excluded from the analysis since he reported an explicit strategy to perform task. The participants sat on a chair, held the handle of a robot manipulandum with their right arm, and made shooting-like quick movements by displacing the handle of the robot on a horizontal plane^24^. A computer projector displayed all visual stimuli onto a flat opaque board that occluded both the manipulandum and the arm. The visuomotor meta-learning paradigm was composed of a sensory-error (S) trial and a monetary feedback (M) trial. In the S trial, the visual cursor was projected on the screen, which provides online feedback of the hand (Figure 1A, left). This cursor was rotated ±7° with respect to the hand movement to induce a sensory prediction error, i.e., the error between the predicted and generated movements. The goal for the shooting was presented with an arc of ±45° instead of a target, and the participants were asked to randomly choose a movement direction and cross somewhere on the arc except the edge, which emphasizes the sensory prediction error and minimizes the extent of how reach error interferes with task performance. This S trial was followed by M trials, in which the small visual target, instead of the arc, was presented, and the participants were asked to shoot at the center of the target as accurately as possible without the cursor feedback. After the shooting movement, the participants received monetary feedback, which was presented as a numerical score (Figure 1A, right). This score was computed as a function of the size of the memory update (learning), which was measured by the aftereffect (i.e., the changes in the reach direction following the previous sensory prediction error). The participants repeated the cycles of one S trial followed by four M trials.

**Figure 1.**
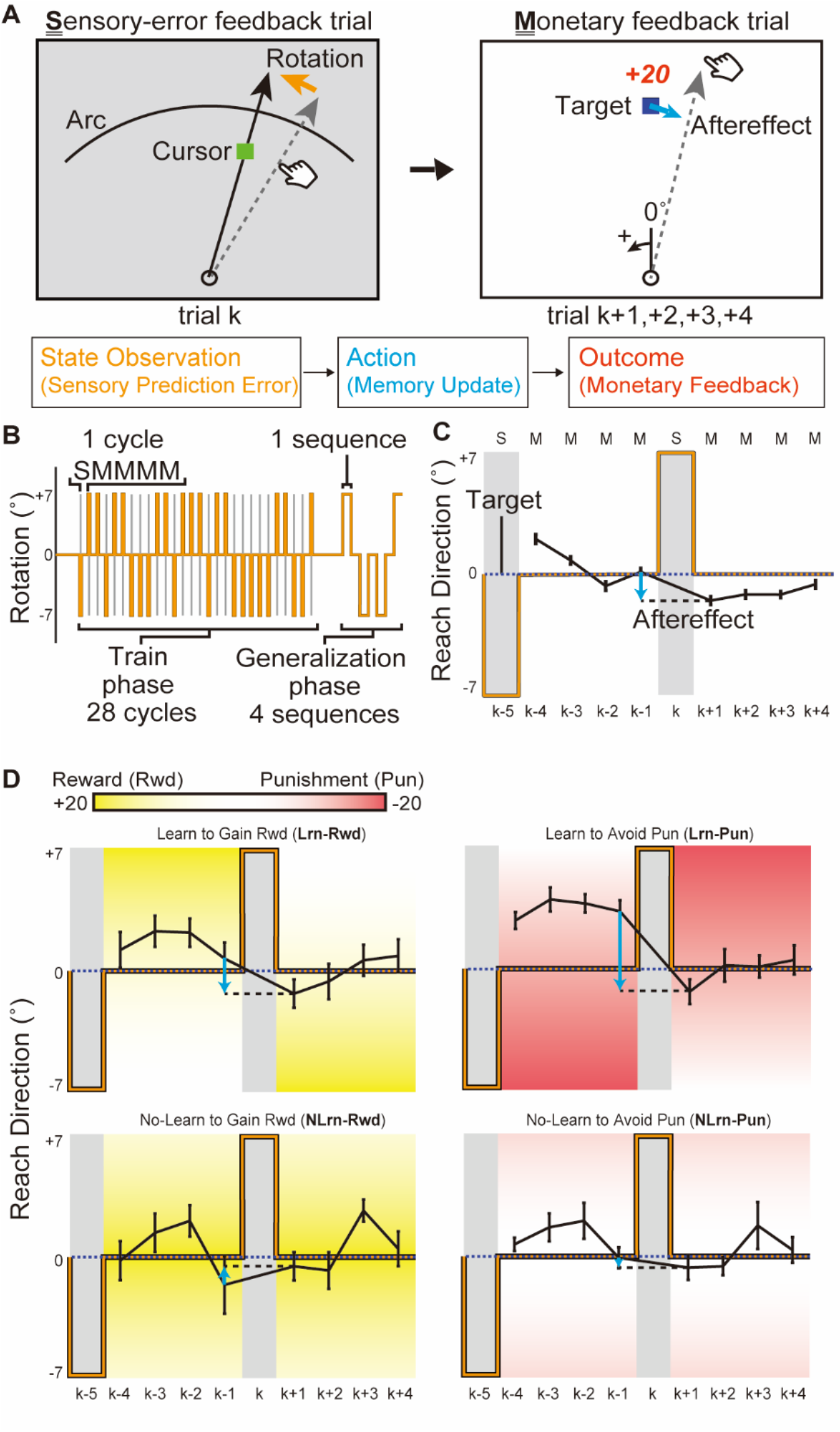
Reward-based visuomotor meta-learning paradigm. (A) Training paradigm. Cursor rotation (orange arrow)-induced sensory prediction error observation (state) in the sensory error (S) trial. The aftereffect size (action, cyan arrow) was scored (outcome) in the monetary feedback (M) trial. (B) One block consisted of the training phase, which included 28 cycles of 1 S trial followed by 4 M trials. After brief washout (null) trials, the generalization phase was composed of four short sequences of visual rotation trials without scores. (C) Plot of reach direction w.r.t. the target direction (memory) for one example rotation pattern in baseline. The aftereffect (cyan arrow) was developed in the opposite direction to the rotation (orange line) in the S (*k*^th^) trial, showing a motor memory update. (D) Memory profile for the same rotation pattern for each group. The scores improved with memory updates toward the yellow zone (large reward) or away from the red zone (large punishment). The error bars indicate standard error of the mean (SEM).

This task structure implements a reinforcement learning problem among trial sequences formalized as a Markov decision process, whereby the agent observes the environmental state, responds according to its policy and is rewarded for the action^28^. In our task, observation of the sensory prediction error (state observation) and the learning rate (policy) yielded a memory update (action), which subsequently generated a monetary feedback (outcome, Figure 1A, bottom). Thus, if the brain learns to maximize the monetary feedback, the action-outcome structure (i.e., how much is gained by how much is learned) should determine the change in learning rate. In addition, if the effect of this action-outcome structure is dominant, the valance which has been considered to influence learning rates^13^ does not influence the change in learning rate.

The experiment contained five blocks, each of which comprised a training phase followed by a generalization phase. The training phase had 28 cycles, each containing one S trial and four M trials. The generalization phase included four short sequences of visuomotor rotation trials to assay the generalizability of meta-learned learning rates to a conventional visuomotor learning task (Figure 1B).

An example of the visual rotation pattern (+7° after −7°) is shown in Figure 1C. In the first (baseline) block, no score was given in M trials. Nevertheless, after the sensory prediction error led by visual cursor rotation was observed at the *k^th^* trial (S trial), the reach direction was updated to compensate for the given rotation. Thus, the change in movement direction at the *k^th^* + 1 trial was significantly different from that at the *k^th^* - 1 trial (cyan arrow in Figure 1C, Wilcoxon paired signed-rank tests, *V* = 726, *p* < 0.00001, *r*=0.70), indicating that the robust aftereffects of memory updates were induced by the sensory prediction errors, which is congruent with previous reports of the roles of sensory prediction errors on updates of motor memory^29^.

In subsequent blocks, the participants were randomly assigned to one of four groups (*n* = 10/group) with different action-outcome structures and valences. In the learn (Lrn) structure, larger aftereffects yielded larger scores, whereas in the not-learn (NLrn) structure, smaller aftereffects yielded larger scores, indicated by the background gradation colors in Figure 1D. The valence determined whether the monetary feedback was positive (reward [Rwd]) or negative (punishment [Pun]). This design required the participants to either learn more to gain more rewards (Lrn-Rwd) and avoid larger punishments (Lrn-Pun) or learn less to gain more rewards (NLrn-Rwd) and avoid larger punishments (NLrn-Pun). To visually observe differences in the regulation of memory updates among these conditions, we focused on the same example rotation pattern (+7° after −7°) as that for the baseline analysis (Figure 1D), and the same Wilcoxon paired signed-rank test was performed. The analysis showed robust aftereffects in the Lrn groups but not in the NLrn groups (Lrn-Rwd: *V* = 53, *p* = 0.0059, *r*=0.80; Lrn-Pun: *V* = 55, *p* = 0.002, *r*=0.91; NLrn-Rwd: *V* = 37, *p* = 0.38, *r*=0.10; NLrn-Pun: *V* = 18, *p* = 0.38, *r*=0.10). That is, when the memory update led to an increase in rewards and a decrease in punishment, the aftereffect was kept robust; in contrast, when the memory update led to a decrease in rewards and an increase in punishment, the aftereffect was attenuated.

We considered that the participants regulated the learning rate in accordance with the action-outcome structure (Lrn v.s. NLrn). To confirm this, for all combinations of perturbations (+7° after +7°, −7° after −7°, +7° after −7°, and +7° after −7°), we estimated the learning rate, *β*, by taking the ratio of the memory update (aftereffect) to the sensory prediction error. In other words, we computed how much the reach direction changed relative to the sensory prediction error in an S trial (see Methods for details). The median *β* was taken for each individual participant and each block for analysis. Figure 2A shows the group mean of individual medians over the blocks for each group and illustrates that the increase or decrease in learning rate over the blocks was different for the groups. To confirm this, we examined the trend of the learning rate over the blocks using a linear mixed-effect model with the action-outcome structure, valence, and block as the fixed effects and participant as the random intercept effect (see Methods for details). This analysis revealed a significant interaction between the action-outcome structure and the block (*t* = −2.65, *p* = 0.009, *R^2^*=0.61). Additionally, we estimated marginal slopes for the change in learning rate over the blocks with respect to the action-outcome structure, which confirmed that the change in learning rate over the blocks was larger in the Lrn groups than in the NLrn groups (*t* = −2.61, *p* = 0.0099). However, we did not find a significant effect of valence (*t* = 1.76, *p* = 0.08). Furthermore, we computed the change in learning rate compared with the baseline for each block and then took the individual participant means of this value across the blocks (Δ*β*, Figure 2B). This result was then analyzed by a reduced linear model with the action-outcome structure and valence as the fixed effects (see Methods for details). Then, we confirmed that the change in learning rate (Δ*β*) was larger in the Lrn group than in the NLrn group (*t* = −2.14, *p* = 0.04, *R^2^*=0.15), while neither valence (*t* = 1.21, *p* = 0.23, *R^2^*=0.15) nor the interaction between action-outcome and valence (*t* = −0.38, *p* = 0.70, *R^2^*=0.15) has a significant effect. These results demonstrate that the participants regulated their learning rates according to the action-outcome structure regardless of valence.

**Figure 2.**
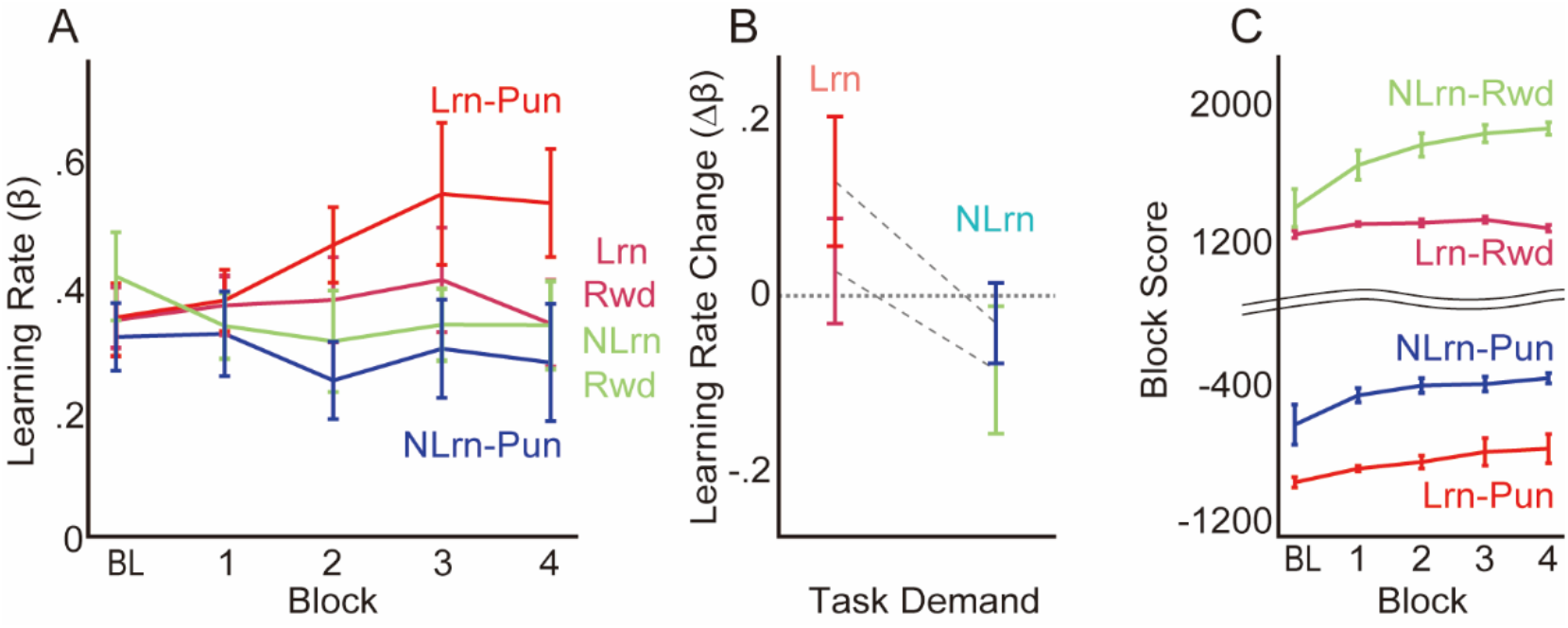
Mean performance scores and learning rates. (A) Estimated changes in learning rates by block. (B) The learning rates changed more in the Lrn group than in the NLrn group. The error bars indicate SEM. (C) Total scores for each block. The baseline (BL) scores were calculated and are shown here.

If these regulations are achieved by reinforcement learning of learning rates, outcomes that were provided as scores should be maximized. Figure 2C shows the across-subjects mean of the block total score in all conditions. Notably, the baseline was not presented to the participants but was calculated in the same way as the following intervention blocks. As a result of the regulation of the learning rate, participants in all conditions increased their monetary outcome (one-sample Wilcoxon-test; Lrn-Rwd: *V* = 48, *p* = 0.04, *r*=0.56, Lrn-Pun: *V* = 49, *p* = 0.03, *r*=0.61 NLrn-Rwd: *V* = 55, *p* = 0.002, *r*=0.91, NLrn-Pun: *V* = 49, *p* = 0.03, *r*=0.61). These results support the hypothesis that the participants both up- and down-regulated the learning speeds via reinforcement meta-learning: instrumental and active meta-learning processes.

Does this reinforcement meta-learning regulate learning rates of motor learning or something else specific for this training task? Subsequently, after the training phase, the participants experienced the generalization phase, which involved a conventional visuomotor rotation task in which both the visual target and the hand cursor were presented and no monetary feedback was provided (Figure 1B). Then, we investigated the learning rate in the generalization phase to determine whether the effect of the action-outcome structure in the reinforcement meta-learning was transferred to a conventional visuomotor rotation task. Additionally, we investigated time-dependent changes within the generalization phase to determine the temporal robustness of the effect. To do so, we estimated the learning rate, *β*, for each individual, block, and sequence, and then the individual mean change from baseline across the blocks, Δ*β*, was calculated for each sequence. Figure 3A shows the group means for individual mean change over the sequences for the Lrn and NLrn groups. The individual mean change in learning rate was analyzed with a linear mixed-effect model with the action-outcome structure, the sequence, their interaction as fixed effects and the participant as the random intercept effect (see Methods for details). The analysis revealed a significant effect of the action-outcome structure (*t* = −2.08, *p* = 0.04, *R^2^*=0.67) and its interaction with the sequence (*t* = 2.58, *p* = 0.01, *R^2^*=0.67). Furthermore, the individual mean changes in learning rate for the training phase correlated with those in the first half (Figure 3B, *R* = 0.43, *t*_38_ = 2.90, *p* = 0.006) but not with those in the second half (*R*=0.25, *t*_38_ = 1.61, *p* = 0.12) of the generalization phase. Notably, although the change in learning rate was greater in the Lrn groups than in the NLrn groups in the first half (vertical axis of Figure 3B, *t* = −2.78, *p* = 0.008, *d* = .88), this phenomenon was not observed for the second half (*t* = −0.74, *p* = 0.46, *d* = .23). These results demonstrate that the regulation of the learning rate by reinforcement meta-learning in the training phase was generalized to the learning rate in motor learning for conventional visuomotor rotation tasks. This suggests that what is updated in the training phase overlaps with the learning rate of visuomotor learning.

**Figure 3.**
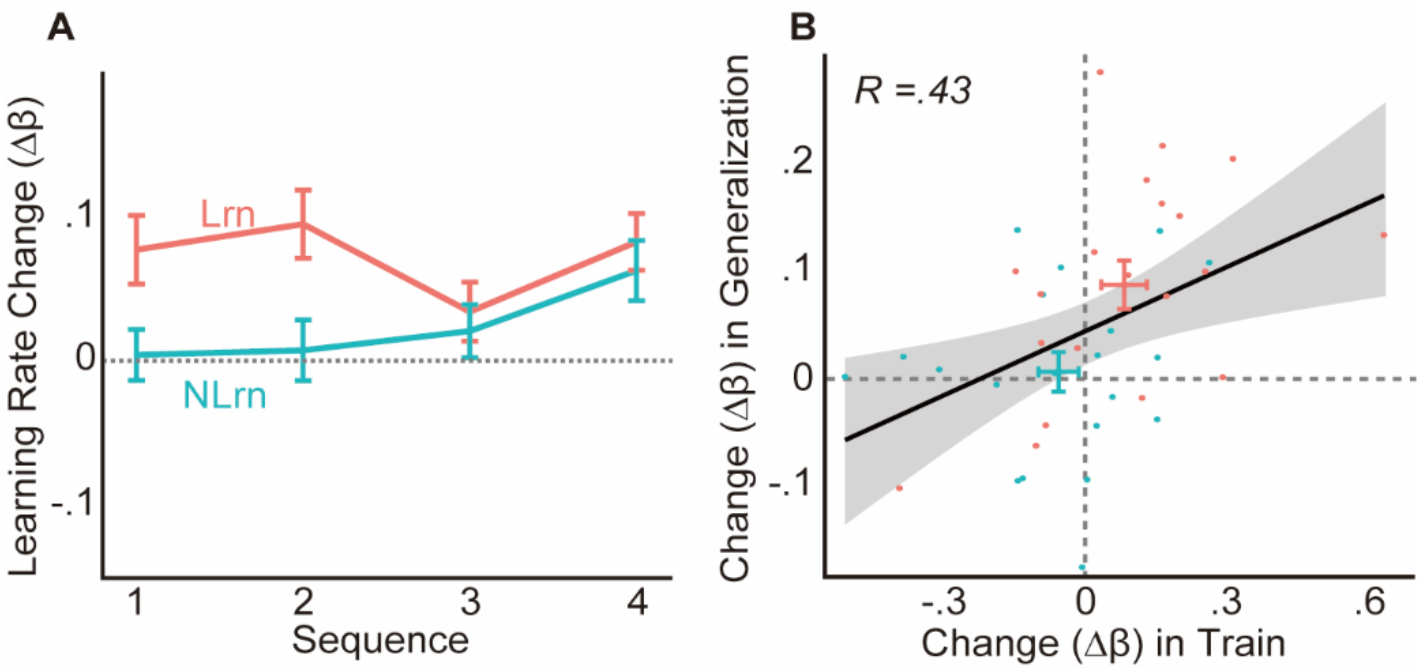
Mean learning-rate changes in the generalization phase. (A) Mean learning-rate changes across the blocks for each sequence. (B) The learning-rate changes in the training phase corresponded with those in the first half of the generalization phase. Each dot represents the data from an individual participant. The error bars indicate SEM.

## Discussion

We found that by presenting rewards as a function of the amount of memory update that was induced by sensory prediction errors, the brain both up- and down-regulated the learning rates to increase rewards as well as to avoid punishment. This effect was gradually developed over the training sessions, and the directionality of regulation (i.e., learn or not-learn) was determined by the action-outcome structure, not by valence. These results suggest that reinforcement learning is employed to regulate learning rates. Furthermore, by examining the generalization of these regulations from the training task to the conventional visuomotor rotation task, we found that these meta-learning effects were transferred to motor learning. This observation indicates that there is an overlapping neural basis between reinforcement meta-learning and visuomotor learning. Thus, our results demonstrate the existence of a reinforcement meta-learning mechanism for motor learning in the human brain.

In machine learning studies, automatic tuning of learning parameters has been a long-standing problem^30^. A computational model of biological reinforcement learning suggests that, in the brain, neuromodulators such as noradrenaline, acetylcholine, and noradrenaline adjust learning parameters of reinforcement learning^31^ via reward-based modulation of these parameters^32^. Recent algorithms of meta-learning in machine learning studies have highlighted a hierarchical structure composed of two reinforcement learning systems: while a low-order reinforcement learning optimizes the weight parameters for action selections in a single learning episode, a high-order reinforcement learning optimizes meta-parameters of the low-order reinforcement learning network to maximize rewards across multiple learning episodes^25,26^. Although the neural implementation of such reinforcement meta-learning was recently discussed^27^, there was no experimental examination. Here, we devised a meta-learning paradigm in which reward feedback for meta-learning, which was provided independently from the sensory error feedback for motor learning, was manipulated as a function of the learning rate of motor learning. Our data demonstrated that while motor memory was updated to minimize the given sensory prediction errors in a single trial, the learning parameter of motor learning was updated to maximize rewards over multiple trials.

The observed meta-learning effect may account for previous reports of learning-rate flexibility during motor learning tasks^6,7,13,33^. For instance, the learning rate increases over the sessions when the perturbation is relatively constant but decreases when the direction of perturbation frequently changes. Importantly, a typical learning paradigm with constant perturbation shares the same action-outcome structure as that for Lrn-Pun, whereby a memory update attenuates movement errors on the next trial, which can be considered avoidance of aversive outcomes^34,35^. In this case, reinforcement meta-learning accelerates learning rates to quickly reduce errors in re-learning (e.g., saving)^6^ or learning with additional punishment signals^13^. Conversely, a non-typical learning paradigm with rapidly changing perturbation shares the same action-outcome structure as NLrn-Pun because the memory update increases errors in the next trial. In this case, reinforcement meta-learning decreases the learning rate. Thus, reinforcement meta-learning explains how the statistics of the perturbation affect learning rates^7,33^.

According to a conventional theory of motor learning, the brain updates motor commands independent of reward to minimize sensory prediction errors^36^. Although motor memory has also recently been found to be updated by rewards, the neural basis of this reward-based motor learning is likely distinct from that of sensory error-based motor learning^24,37^. According to these previous studies, rewards might not interact with sensory prediction errors during motor learning. Here, our data demonstrate that the sensory prediction errors and the rewards presented in separated trials were integrated at the higher-level motor learning system, i.e., meta-learning, to regulate learning rates.

In theory, to establish reinforcement meta-learning for motor learning, the sensitivity to sensory prediction errors should be evaluated by the rewards. Thus, our results suggest a close interaction between rewards and sensory prediction errors during motor learning. Because research evidence suggests the involvement of cortico-basal ganglia networks in reinforcement meta-learning^27^ and the cerebellum in sensory error-based learning^38,39^, reinforcement meta-learning for motor learning is likely mediated by the functional connectivity between these two learning systems. Thus, the anatomical projections between the basal ganglia and the cerebellum could have a computational role in reinforcement meta-learning^40^. This possibility is further supported by recently reported reward-related signals in cerebellar inputs^41^ and outputs^42^ during motor control tasks. We suggest that these interactions between the basal ganglia and the cerebellum play a key role in optimizing learning parameters for motor learning via reinforcement learning.

## Acknowledgments

This work was supported by KAKENHI (grant numbers 17H06023 and 19H04977). T.S. is grateful for financial support from the JSPS Research Fellowship for Young Scientist and KAKENHI (19J20366).

## Author Contributions

Conceptualization, T.S. and J.I.; methodology, T.S. and J.I.; investigation, T.S. and J.I.; formal analysis, T.S. and N.S.; writing – original draft, T.S., and J.I.; writing – review & editing, T.S., N.S., and J.I.; funding acquisition, J.I.

## Methods

### Participants

Forty one right-handed participants (25 males; aged 19–37 years, *μ* = 24) volunteered for this experiment. Their handedness was confirmed by the Edinburgh Handedness Inventory, and they reported no history of neurological or motor disorders. We excluded one subject who reported an explicit strategy to perform the task from the analysis. They were paid 1,640 JPY for their participation, with an additional performance-based compensation up to 1,000 JPY.

### Task design

#### General

Participants performed the task using a robot manipulandum^24^ that moved only in the horizontal plane. They sat on a chair and held the robot handle in their right hand. A horizontal, flat, opaque board covered the task space, occluding the hand and forearm. A computer projector was fixed above and displayed visual information on the board.

In each trial, participants made a rapid shooting movement when a target appeared 10 cm away from the starting point. To control for use-dependent learning^43^, the target location was pseudo-randomly selected from 1 of 7 locations: −15°, −10°, −5°, 0° (directly in front of the participant), 5°, 10°, and 15°. The counter-clockwise direction was defined as positive in the angular coordinates (**Figure 1A**). To maintain similar kinematics across trials, “Too Fast” or “Too Slow” was displayed as a warning when the movement duration was <200 ms or >300 ms, respectively.

#### Trial type and task schedule

There were four trial types: Null, Sensory-error feedback (S), Monetary feedback (M), and Generalization. In Null trials, participants made shooting movements toward the targets with veridical online cursor feedback. In S trials, the targets were replaced with an arc spanning ±45°, centered on the home position and with 10 cm radius (**Figure 1A**)^44^. Participants were asked to cross the arc while trying to distribute their reach direction across S trials. Online cursor feedback was rotated ±7° from hand movements to induce errors between the predicted hand position and the visual feedback. The rotation direction was pseudo-randomly selected (**Figure 1B**). Note that the use of an arc instead of a target minimizes the extent of how reach errors interfere with task performance and, thus, emphasizes errors between the observed and predicted hand movements, i.e., sensory predictor errors. In M trials, participants made shooting movements toward targets without online cursor feedback. Upon movement completion, monetary feedback was presented as a numerical score above the target (**Figure 1A**). Scores were computed based on the reach direction from the target, with different computations across the experimental conditions (see below). Finally, Generalization trials were identical to Null trials except that, similar to S trials, online cursor feedback was rotated ±7° from the hand movement, as in conventional visuomotor rotation tasks^45^.

Participants performed first one Baseline block, in which no monetary feedback was provided, and then four blocks with monetary feedback. Each block comprised a Train phase with 28 cycles of 1 S trial and 4 M trials, followed by a Generalization phase. Brief washout blocks of 14 Null trials were inserted before each phase. The Generalization phase comprised 4 Sequences of 5 consecutive cursor rotation trials, with 5 Null trials between each Sequence (**Figure 1B**). The aim of the Generalization phase was to test whether the participants generalized the meta-learning effect formed in the Train phase to a conventional visuomotor learning task.

### Estimation of learning rates in the Train phase

We estimated the learning rates in the Train phase with a simple first order state-space model of memory updates to track transitions in reach direction in response to the experienced cursor rotation, as in previous work^7^. In this framework, motor learning is considered a process that estimates perturbations imposed in the task environment. Specifically in the visuomotor rotation task, the executed motor plan *u*^(*t*)^ at trial *t* determined the direction of the hand movement *h*^(*t*)^,

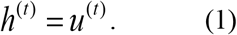

While the hand movement was not directly observable for participants, a cursor projected on the screen provided online feedback of the hand motion while the visual rotation *p*^(*t*)^ was imposed between the hand movement and the visual cursor:

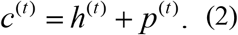

The brain may predict hand movement direction *ĥ*^(*t*)^ based on the estimation of the perturbation *x*^(*t*)^ and the efference copy of the motor plan *u*^(*t*)^:

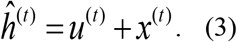

To minimize the sensory prediction error *c*^(*t*)^ − *ĥ*^(*t*)^, the brain updates the estimate of the perturbation with the following learning rule:

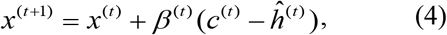

where the learning rate *β* characterizes the rate of learning.

Here, we estimated the learning rate *β* in the Train phase using the data of hand directions in a triplet of M, S, and M trials, following previously developped methods^46^. That is, considering a certain trial of S trials at the trial *t* = *k*, *β*^(*k*)^ was estimated using the measured hand movement direction *h*^(*k*)^ and the given cursor rotation *p*^(*k*)^ in S trials at *t* = *k*, *h*^(*k*−1)^ and the presented target direction *T*^(*k*−1)^ in M trials at *t* = *k* - 1, as well as *h*^(*k*+1)^ and *T*^(*k*+1)^ in M trials at *t* = *k* + 1. Because in M trials, the visual target *T* was presented and the participants were explicitly instructed to cross the target with their unseen but estimated hand position, the participants’ estimation of their hand direction should closely match the target direction. We thus assumed *T*^(*k*−1)^ = *ĥ*^(*k*−1)^ and *T*^(*k*+1)^ = *ĥ*^(*k*+1)^. From (1) and (3), we have *x*^(*k*−1)^ = *T*^(*k*−1)^ − *h*^(*k*−1)^ and *x*^(*k*+1)^ = *T*^(*k*+1)^ − *h*^(*k*+1)^. Importantly, because the cursor was not given to participants in M trials at *t* = *k* - 1, the sensory prediction error *c*^(*k*−1)^ − *ĥ*^(*k*−1)^ was absent and thus, no memory update was engaged. Hence, we assumed *x*^(*k*)^ = *x*^(*k*−1)^. For subsequent S trials at *t* = *k*, because the sensory prediction error *c*^(*k*)^ − *ĥ*^(*k*)^ was present, the memory was updated in accordance with (4).

Using (4), we estimate *β*^(*k*)^ by:

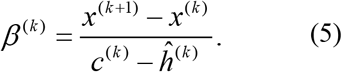

According to (1), (2), (3), and (5), we have:

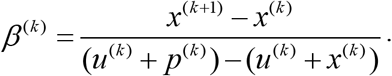

Then, by applying *x*(*k*) = *x*^(*k*−1)^, we have:

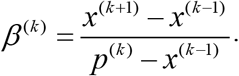

Finally, by substituting *x*^(*k*−1)^ by *T*^(*k*−1)^ − *h*^(*k*−1)^ and *x*^(*k*+1)^ by *T*^(*k*+1)^ − *h*^(*k*+1)^, we have

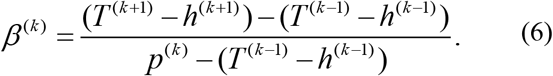

Because we measured *T*^(*k*+1)^ − *h*^(*k*+1)^ and *T*^(*k*−1)^ − *h*^(*k*−1)^ in M trials and manipulated *p*^(*k*)^ as a cursor rotation in S trials, *β*^(*k*)^ can be estimated for each triplet of M-S-M trials in the Train phase. Note that the denominator *p*^(*k*)^ − (*T*^(*k*)^ − *h*^(*k*+1)^) occasionally approaches 0, resulting in an unreliable estimation of *β*. To prevent this, we used the median instead of the mean for each Train phase for each participant before performing statistical analyses.

A potential limitation of the above method is that it does not account for the effect of another type of memory that is updated by reward signals^24^, which might be formed in M trials at *t* = *k* - 1. If this reward-based motor memory exists, it affects the measured reach direction in subsequent S trials at *t* = *k* and M trials at *t* = *k* + 1. This could potentially bias the approximation of the sensory prediction error *p*^(*k*)^ − *x*^(*k*−1)^ by *p*^(*k*)^ −(*T*^(*k*−1)^ − *h*^(*k*−1)^) and thus, bias the estimation of *β*. However, according to the theory of reward-based motor memory^24^, the directions of the bias is opposite in the two cases of perturbation sequences: whether the direction of the perturbation switches or remains the same. Because the perturbation sequence was pseudorandomized, this bias should be predominantly eliminated when we calculate the median over each block. If a small effect of this bias remains, it would likely lead to an underestimation of *β*for Lrn groups and overestimation for NLrn groups; however, this was not observed in our data. As a final analysis of this potential confounding factor, we examined the generalization of the meta-learning in the Train phase to a conventional visuomotor rotation task where no monetary feedback was given.

### Estimation of learning rates in the Generalization phase

Using the state-space model of the memory update (1)–(4), the learning rate was estimated via least square to fit each subject’s data of measured hand directions. In the Generalization phase, we assumed that the participants aimed at the presented target *T*^(*t*)^ with their prediction of the hand direction *ĥ*^(*t*)^, so that *ĥ*^(*t*)^ closely matched with *T*^(*t*)^ (i.e, *ĥ*^(*t*)^ = *T*^(*t*)^). They observed the error between the cursor and the prediction of the hand *c*^(*t*)^ - *ĥ*^(*t*)^. Thus, the state-space model of the memory update (4) provides us the simulated sequences of *x*^(*t*)^ over trials using the measured sequence of *c*^(*t*)^ − *T*^(*t*)^ for a given *β*. The model also generates simulated sequence of participants’ reach error *E_S_*^(*t*)^ = *T*^(*t*)^ − *h*^(*t*)^ in accordance with (1) and (3). In the experiment, we measured the actual participants’ reach error *E_M_*^(*t*)^ = *T*^(*t*)^ − *h*^(*t*)^. Thus, following a previous method^4^, we estimated the learning rate *β* that minimized the sum of the least square error between *E_S_*^(*t*)^ and *E_M_*^(*t*)^ over each step-perturbation sequence of the three Null trials and the five cursor rotation trials.

### Experiment groups and score calculations

There were two independent variables, Action-Outcome structure and Valence, respectively, with two levels for each. This resulted in four experiment groups (**Figure 1D**). The Action-Outcome structure determined how the monetary feedback was computed in four consecutive M trials (*t* = *k* + *i*,*i* = {1,2,3,4}) after the S trial (*t* = *k*) both as a function of the memory *x*^*k*+*i*^, measured as a reaching angle with respect to the target *T*^(*k*+*i*)^ − *h*^(*k*+*i*)^, and as a function of the given perturbation *p*^(*k*)^. This regulated whether the memory updates in response to the sensory prediction error *β*^(*k*)^(*c*^(*k*)^ − *ĥ*^(*k*)^) were encouraged (Lrn) or discouraged (NLrn). Specifically, for Lrn, larger aftereffects of the exposure to the cursor rotation corresponded to larger scores, whereas, for NLrn, smaller aftereffects corresponded to larger scores. In contrast, Valence determined whether monetary feedback was positive (i.e., reward) or negative (i.e., punishment). Therefore, participants could learn more from sensory prediction error to gain more rewards (Lrn-Rwd) or to avoid larger punishments (Lrn-Pun), allowing them to improve their scores with greater memory updates. Alternatively, they could learn less from sensory prediction error to gain more rewards (NLrn-Rwd) or to avoid larger punishments (Lrn-Pun), allowing them to improve their score with less memory updates.

The score ranges for the Rwd and Pun groups were set to [0,20] and [-20,0], respectively. The Lrn-Rwd/Pun group earned the highest score by showing 100% or more memory for the last experienced rotation in the previous S trial (i.e., 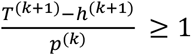 in *i*^th^ M trial within the same cycle), and the score was reduced by 1 point for every 10% less memory until the lowest score (i.e., −100% learning) was reached. The Lrn-Rwd/Pun group earned the highest score by showing 0% memory, (i.e. *T*^(*k*+*i*)^ − *h*^(*k*+*i*)^ = 0), and the score was reduced by 1 point for every 10% more/less memory until the lowest score was reached. These were represented as background color patterns in **Figure 1D**.

### Instructions

Before the task, participants were instructed about the experimental flow for the Train phase and the stimuli and feedback in the S and M trials. They were also explicitly informed that the total score would determine their additional monetary compensation and that the task goal was to maximize their compensation by crossing the target with their hand as closely as possible. In the Rwd conditions, they were told that additional compensation was initially minimum (0 JPY) and accumulated throughout the task. In the Pun condition, participants were told that additional compensation was initially maximum (1,000 JPY) and subtracted throughout the task. In addition, in the S trial, they were told to vary movement directions from trial to trial. This was to avoid the formation of use-dependent behavior^43^. In addition, they were not informed of visual rotation or the relationship between their reach direction and score size to prevent the use of cognitive strategy^47^. Their unawareness of visual rotation throughout the task was confirmed by a written questionnaire after the task was completed asking if they felt a discrepancy between the hand and the cursor during movements.

### Statistical analyses

#### Removal of target location-dependent bias

Due to a gap in height between the physical hand position and the cursor projected on the screen, a bias between the target direction and the reach direction was inevitable. To remove this bias from the analysis, we calculated the mean reach direction for each target location in the Baseline and subtracted from it in the Train phase for all blocks.

#### Memory update following a specific visual rotation

In **Figure 1C,** individual mean reach directions with respect to target direction, *T* − *h*, in the *k*-1^th^ and *k*+1^th^ trials were compared by Wilcoxon paired signed-rank test to examine the aftereffect (i.e., if the reach direction changed after the participants experienced a sensory prediction error in the *k*^th^ trial). Because there was no difference across groups in Baseline, all individual data were combined (*n* = 40). The same test was performed for memory updates in the other four blocks on individual mean differences of *T* − *h* in the *k*-1^th^ and *k*+1^th^ trials across the blocks for each group (**Figure 1D**, *n* = 10/group).

#### Learning rate in the Train phase

We first used a linear mixed effect model (LMEM) with Action-Outcome structure, Valence, and Block as the fixed effects and Participant as the random intercept effect to examine the change in *β* over the course of the task, as shown in **Figure 2A**. All interactions between the fixed effects were included. In R with lme4 package^48^, the model was written as: *lmer*(*β* ~ *Action-Outcome* * *Valence* * *Block* + (1|*Participant*)). In addition, to estimate the marginal slopes of the learning rate change over blocks with respect to Action-Outcome or Valence, we then developed a reduced model without interactions between Action-Outcome and Valence. The model was written as: *lmer* (*β* ~ (*Action-Outcome* + *Valence*) * *Block* + (1|*Participant*)).

Then, to examine the overall change in learning rate across the task, the change in *β* from Baseline (Δ*β*) was calculated for each block and then averaged across blocks for each participant and fit with the following model: *lm*(Δ*β* ~ *Action-Outcome* * *Valence*). Note that because one data point was obtained for each participant, no random effects were included in this model.

#### Block total score performance

To evaluate whether score performance improved, we calculated the total score for each block including Baseline where the score was calculated in the same manner without being presented to the participants, as shown in **Figure 2C**. Then, the change from Baseline was calculated and averaged across blocks for each participant. The change for each group was analyzed by one-sample Wilcoxon signed-rank test with a hypothetical mean of 0 (*n* = 10/group).

#### Learning rate in the Generalization phase

The change in learning rate Δ*β* in the Generalization phase was analyzed similarly to that in the Train phase. Again, individual mean change from Baseline across blocks was calculated by subtracting the Baseline average *β* across Sequence from *β* in the subsequent blocks. Then, Δ*β* was analyzed with a LMEM, with Action-Outcome structure, Sequence, and their interaction, as fixed effects and Participant as random effect intercepts: *lmer*(Δ*β* ~ *Action-Outcome* * *Sequence* + (1|*Participant*)). Note that Valence was removed from the model based on the analysis of the change in *β* in the Train phase, which failed to show its significant effect.

In addition, to examine time-dependent changes within the Generalization phase more closely, the data were separated into the first and second half of the phase. To examine if the change in each half of the Generalization phase reflected the change in the Train phase, a Pearson correlation analysis was then performed on the changes between the Train and the first/second half of Generalization phases. Finally, Δ*β* from each half were analyzed with a linear model with Action-Outcome as the only factor.

#### Statistical package, coding, and validation of linearity

All data processing and statistical analyses were performed in R version 3.5.1 using the lm, lme4^48^, lmerTest^49^, and emmeans^50^ packages. For the linear model analyses, effect coding was used to represent the categorical variables (i.e., Action-Outcome and Valence), except in estimation of marginal slopes where dummy coding was used, and the numerical variables (i.e., Block and Sequence) were represented in z-score (i.e., centered and normalized). Lrn and Rwd were set as reference in coding (i.e., negative value in effect coding and zero in dummy coding). The assumption of linearity was validated by (1) confirming by log-likelihood ratio test on BIC that a model that treats numerical variables as categorical does not produce significantly better fit than a model that treats them as continuous, and (2) confirming the normality of residuals by Shapiro-Wilk test.

